## Supplemental data for "DENV-specific IgA contributes protective and non-pathologic function during antibody-dependent enhancement of DENV infection"

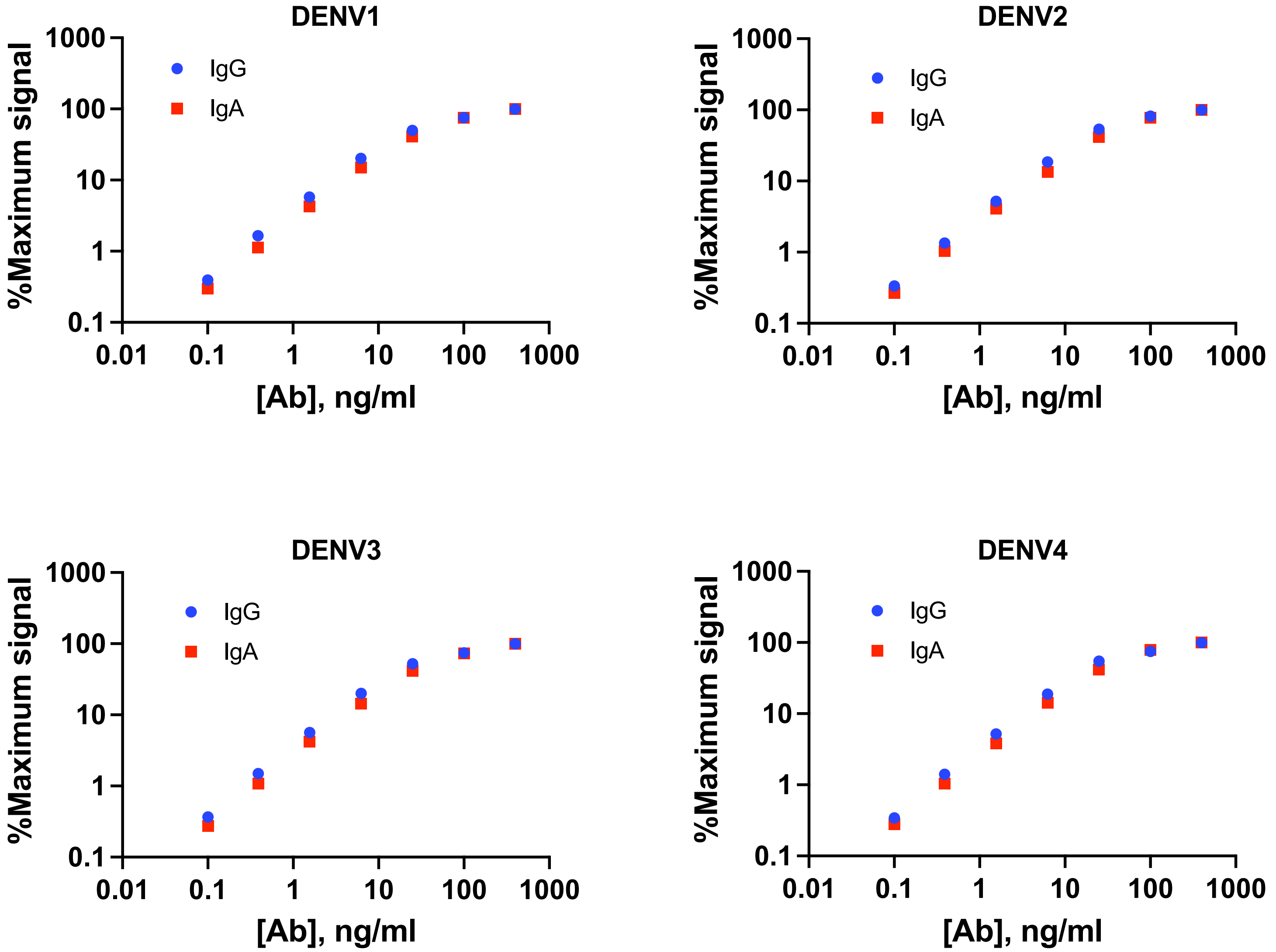


**Supplemental Figure 1.** Assessment of DENV1-4 E binding of VDB33-IgG and VDB33-IgA

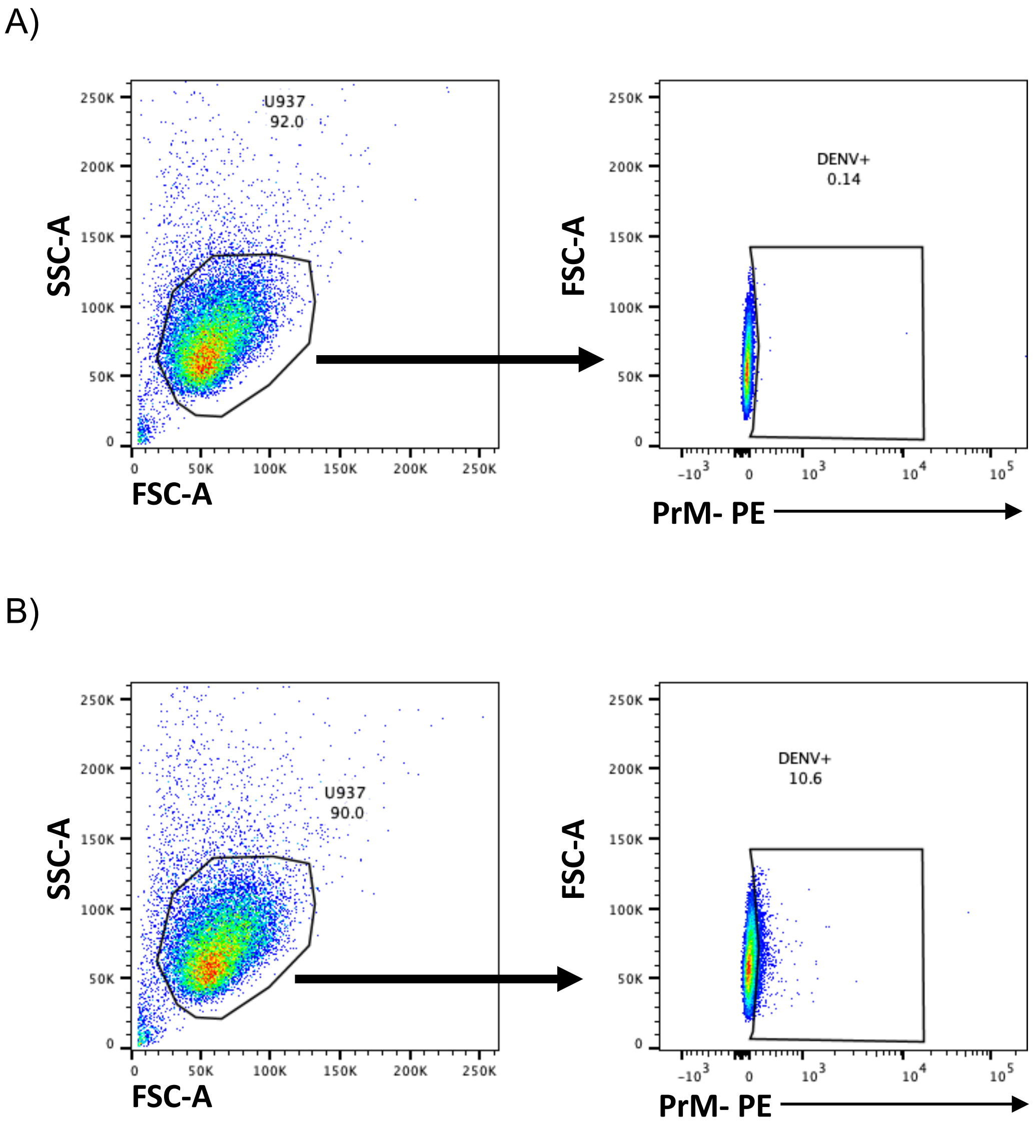


**Supplemental Figure 2:** gating strategy and representative flow plots for **A)** uninfected and **B)** antibody-enhanced infection of U937 cultures

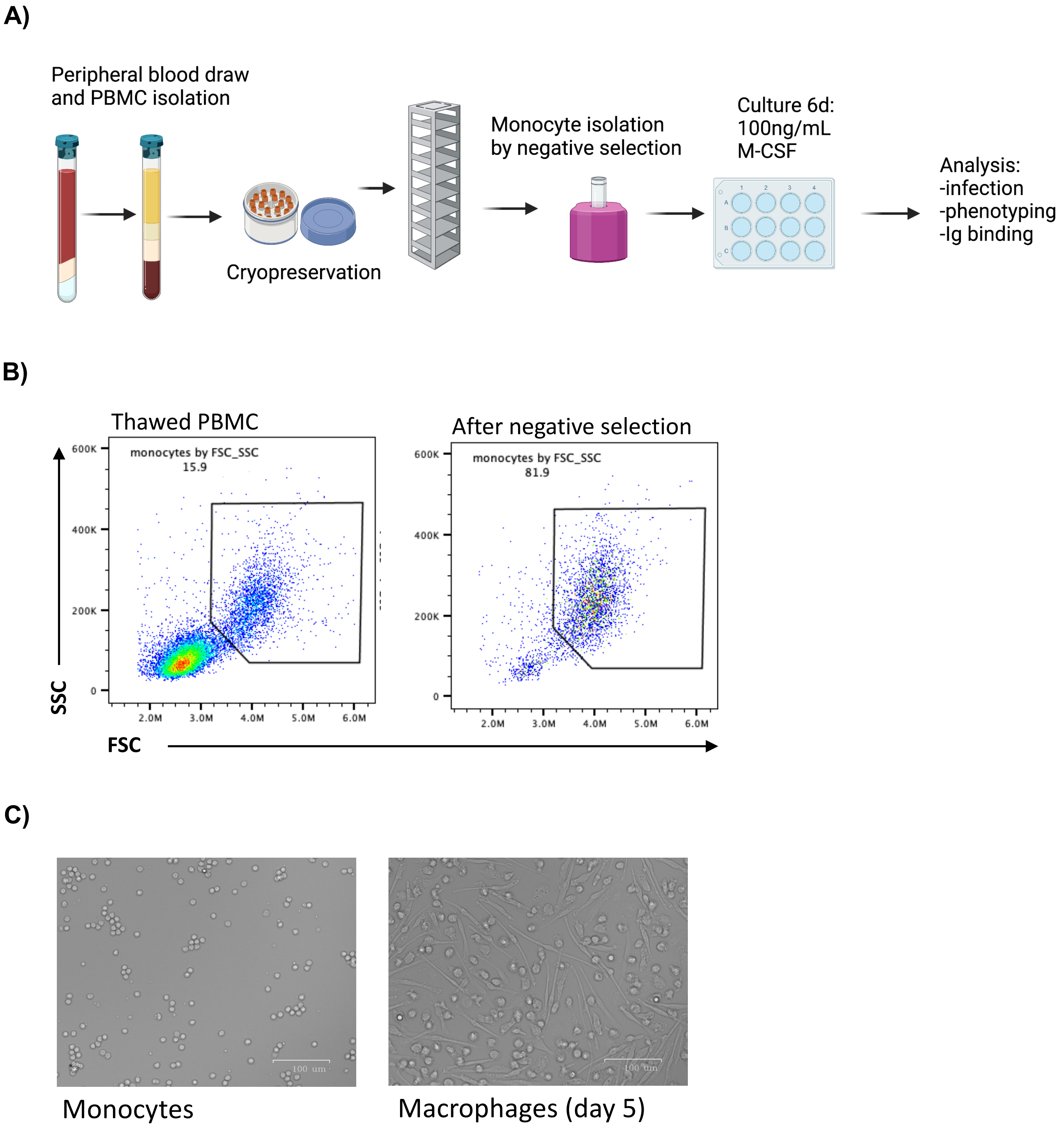


**Supplemental Figure 3: A)** workflow for isolating and cryopreserving human PBMC, isolating monocytes, and differentiation into macrophages. **B)** Flow cytometry analysis of thawed PBMC before and after negative selection for monocytes. **C)** Light micrograph of freshly plated monocytes (day 1) and M-CSF differentiated macrophages (day 5)

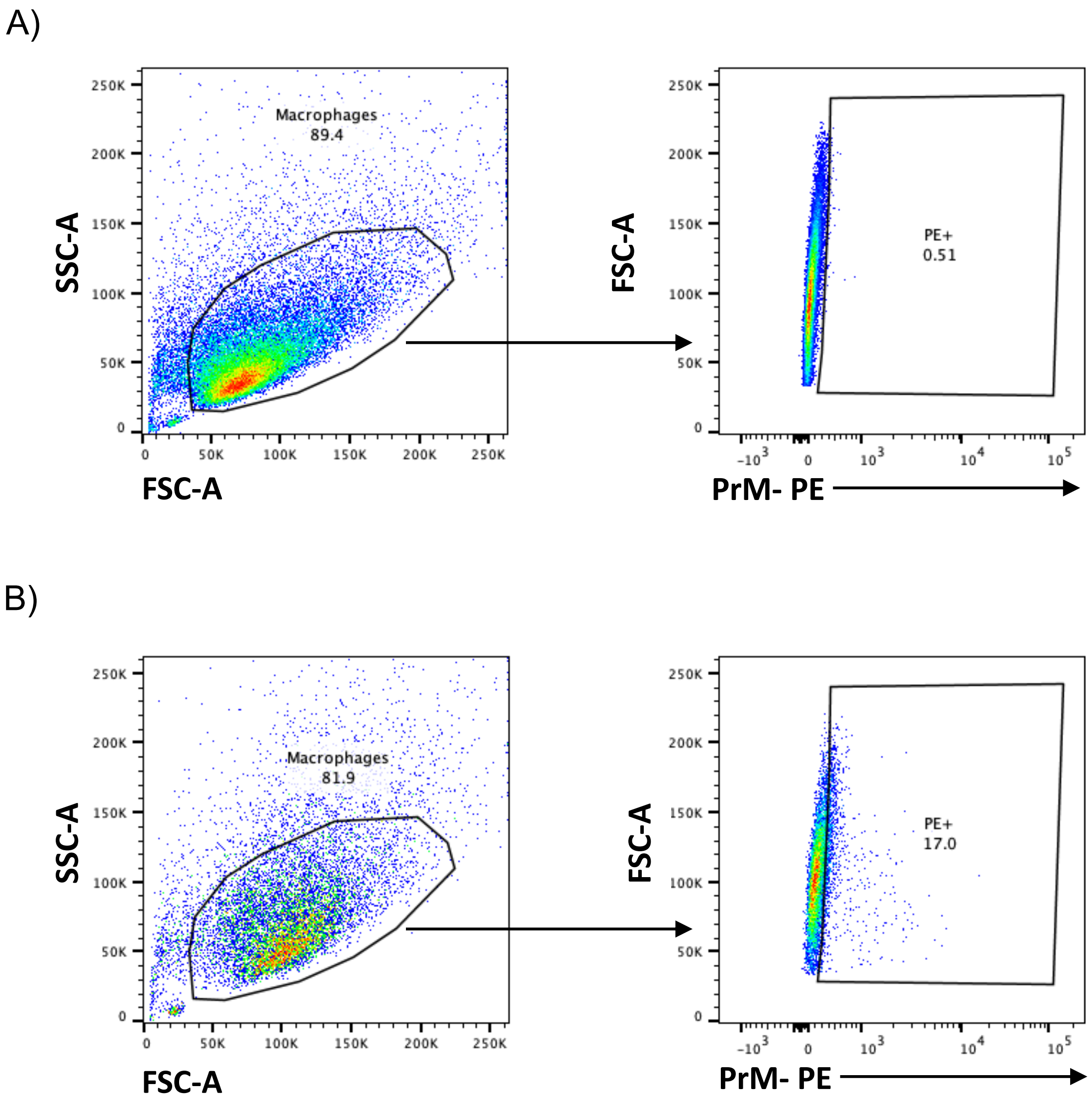


**Supplemental Figure 4:** gating strategy and representative flow plots for **A)** uninfected and **B)** antibody-enhanced infection of macrophage cultures

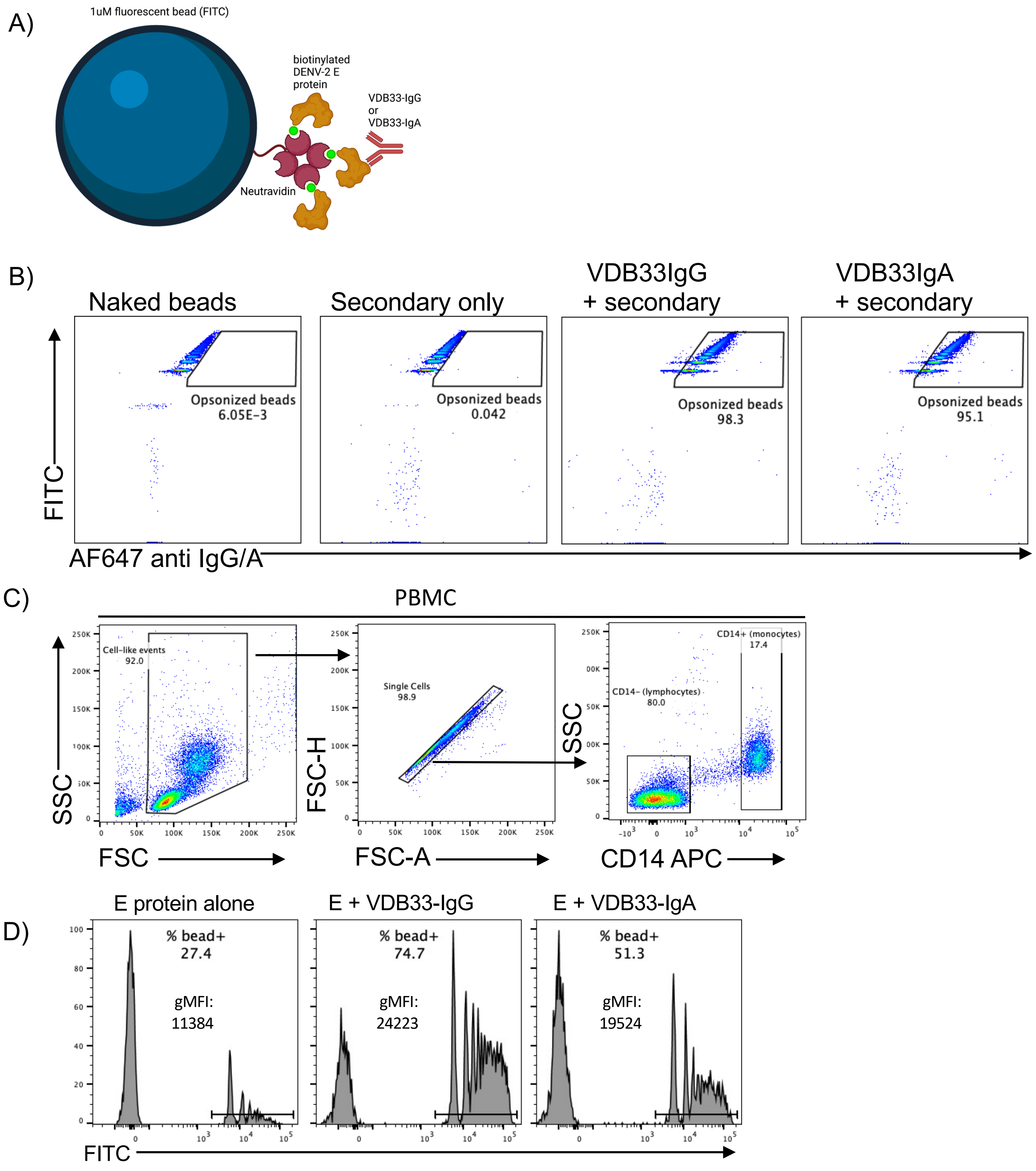


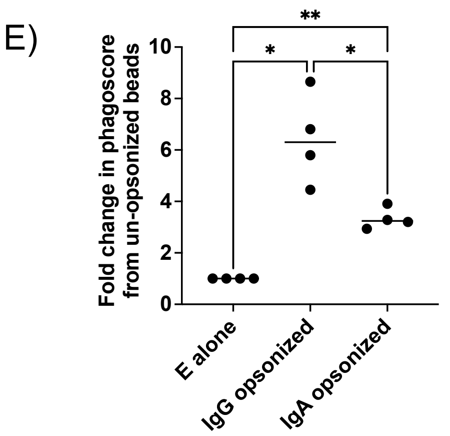


**Supplemental Figure 5: IgG-more efficiently opsonizes polystyrene beads for phagocytosis than IgA.** A) Diagram of DENV E-protein conjugated bead complexes. B) Flow cytometry plots showing opsonization of E-conjugated beads by VDB33-IgG or –IgA. C) Gating strategy for PBMC incubated with fluorescent beads. For final analyses, cells were gated on single live cells, CD14+. D) Representative histograms of the CD14+ monocyte population, showing a sub-population which has internalized fluorescent beads. Labeled E) Plot of phagoscores for 4 biological replicates, normalized to scores of monocyte uptake of un-opsonized beads. Phagoscores are calculated as the product of the % of monocytes positive for bead fluorescence multiplied by the gMFI of the positive population.

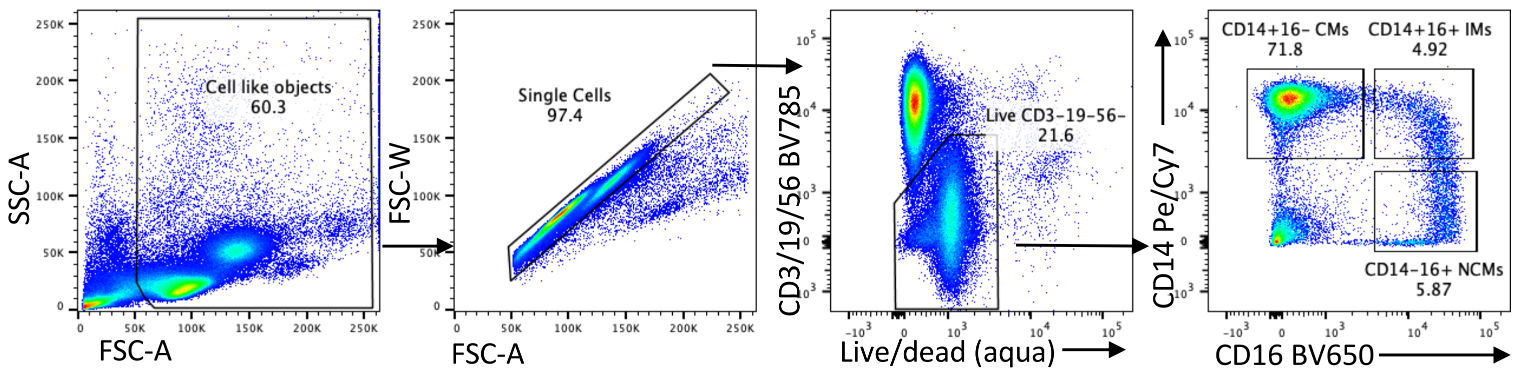


**Supplemental Figure 6:** Gating strategy for analysis of FcR expression on DHIM PBMC

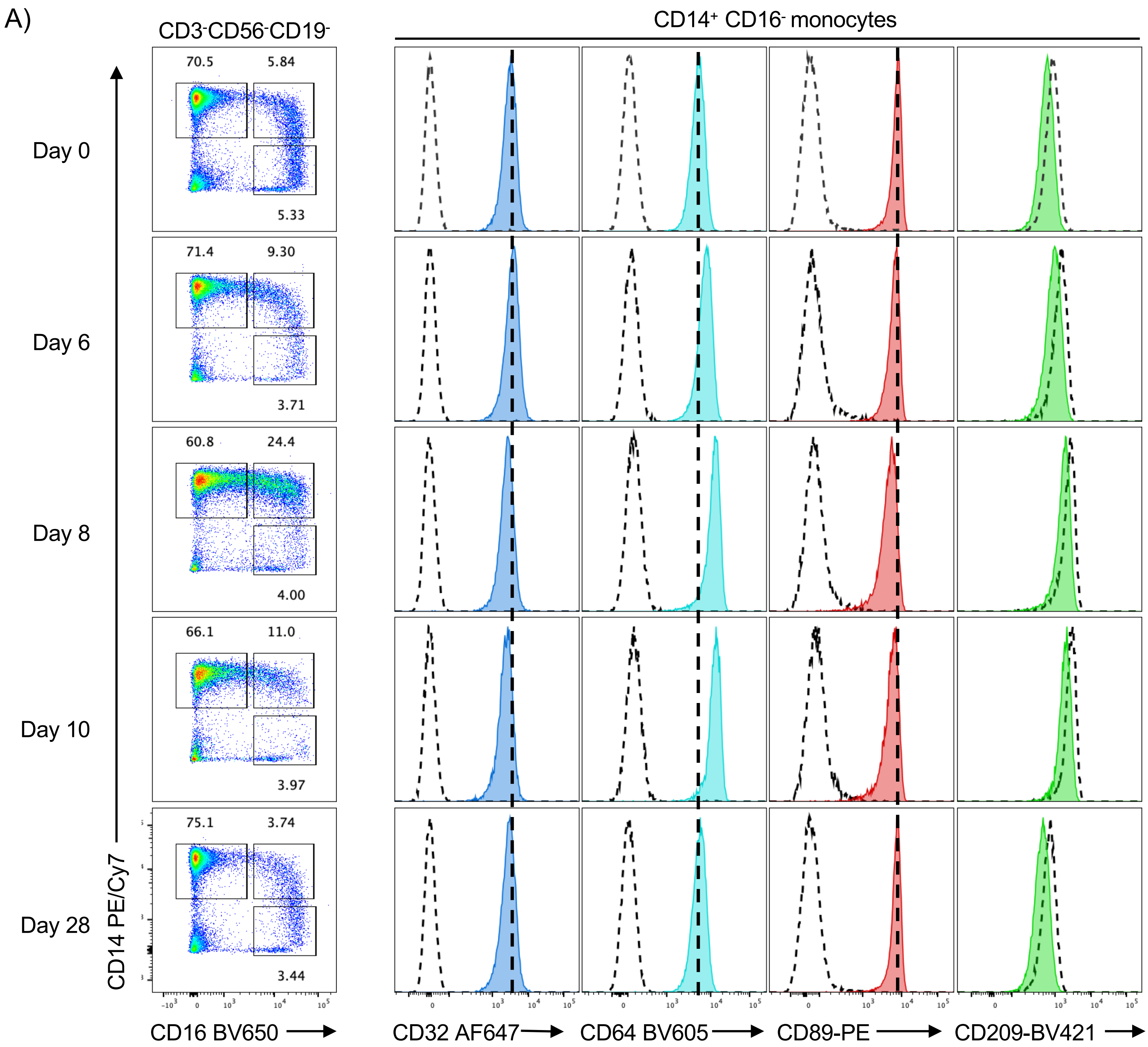


**Supplemental Figure 7: Evolution of monocyte FcR expression over the course of dengue infection.** Cryopreserved PBMC from a DENV-3 infected patient at the indicated timepoints were analyzed by flow cytometry. **A)** Gated on live CD3-19-56-14+16- (classical) monocytes.

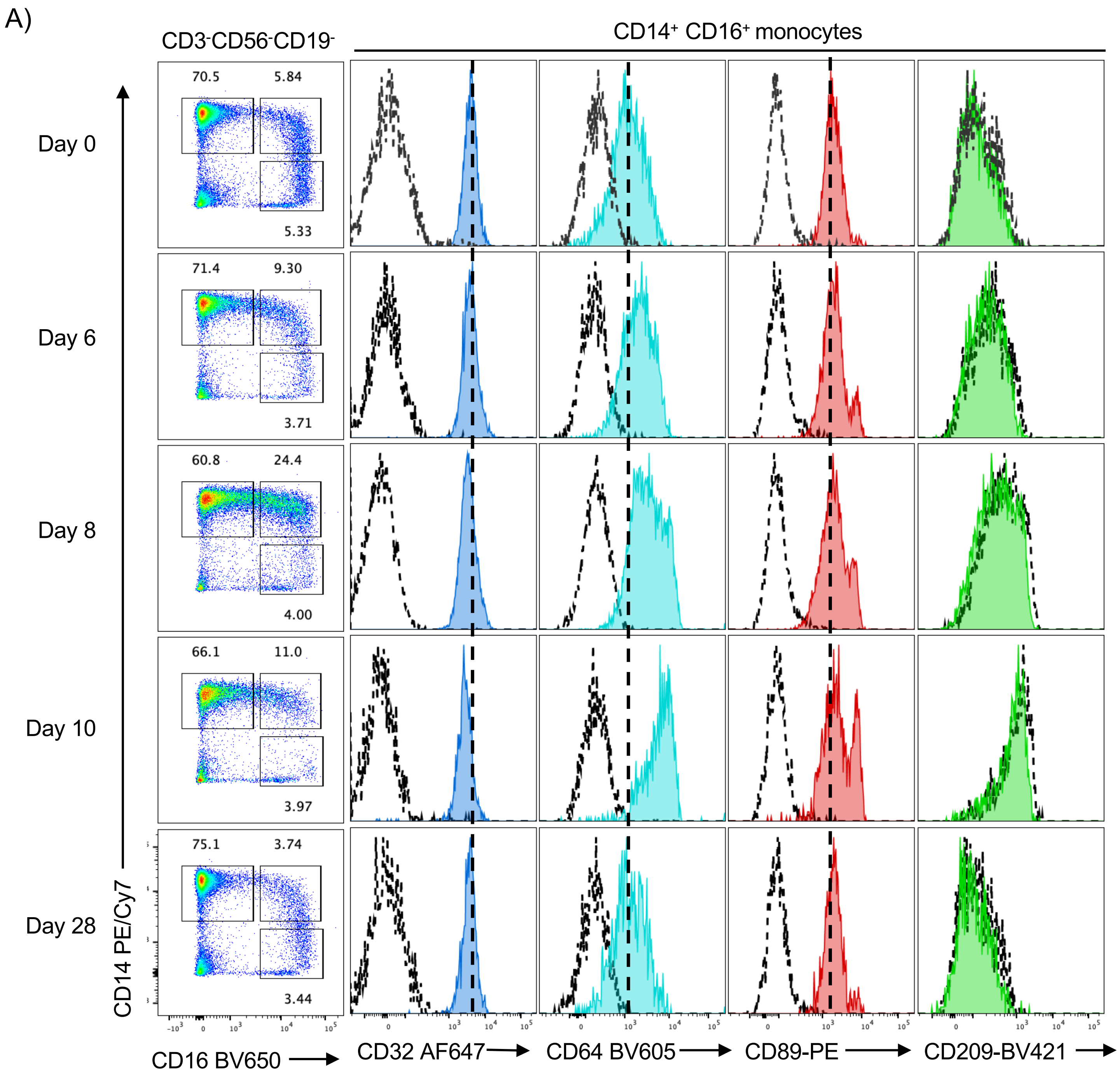


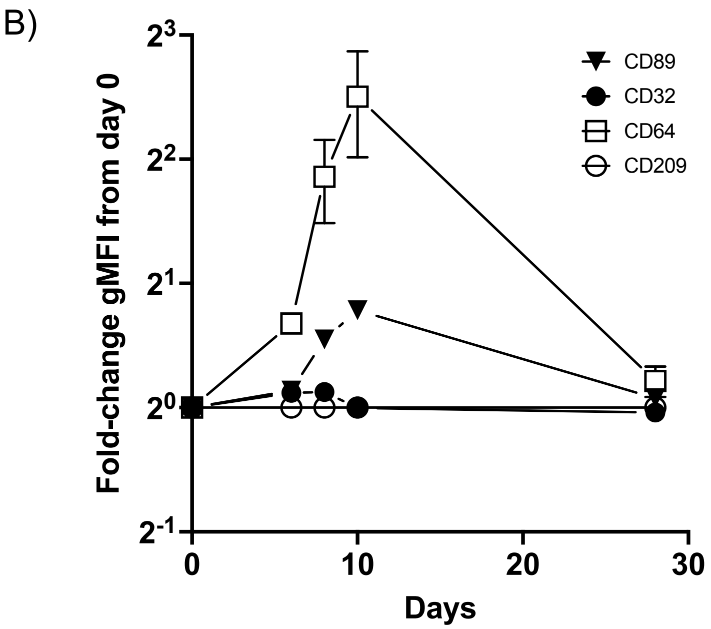


**Supplemental Figure 8: Evolution of monocyte FcR expression over the course of dengue infection.** Cryopreserved PBMC from a DENV-3 infected patient at the indicated timepoints were analyzed by flow cytometry. **A)** Gated on live CD3-19-56-14+16+ (intermediate) monocytes. **B)** Plotted is fold-change of geometric mean fluorescence intensity relative to day 0. Stained samples were isotype control-subtracted prior to fold-change analysis

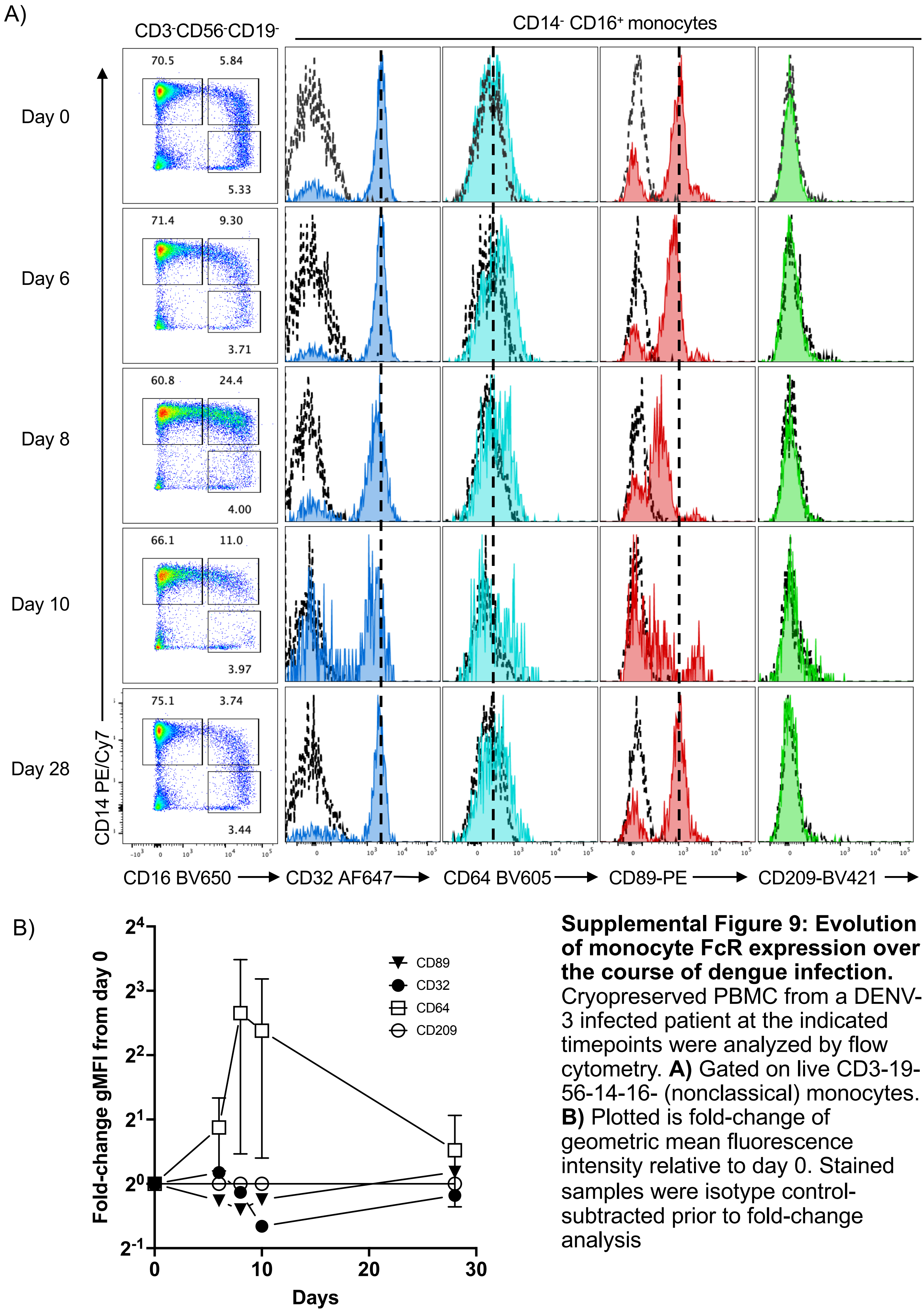


**Supplemental Figure 9: Evolution of monocyte FcR expression over the course of dengue infection.** Cryopreserved PBMC from a DENV-3 infected patient at the indicated timepoints were analyzed by flow cytometry. **A)** Gated on live CD3-19-56-14-16- (nonclassical) monocytes. **B)** Plotted is fold-change of geometric mean fluorescence intensity relative to day 0. Stained samples were isotype control-subtracted prior to fold-change analysis

**Supplemental Table 1.** Antibodies used for flow cytometry

| Antibody | Clone | Manufacturer | Catalog # | Lot # | Dilution |
|---|---|---|---|---|---|
| CD14 PE-Cy7 | 63D3 | Biolegend | 367111 | B371303 | 1:80 |
| CD89 PE | A59 | Biolegend | 354103 | B313937 | 1:160 |
| CD32 AF647 | FUN-2 | Biolegend | 30312 | B378968 | 1:160 |
| CD3 BV785 | OKT3 | Biolegend | 317329 | B360623 | 1:320 |
| CD19 BV785 | HIB19 | Biolegend | 302239 | B355937 | 1:40 |
| CD56 BV785 | 5.1H11 | Biolegend | 362549 | B350905 | 1:40 |
| CD16 BV650 | 3G8 | Biolegend | 302041 | B272555 | 1:320 |
| CD64 BV605 | 10.1 | Biolegend | 305033 | B326358 | 1:80 |
| CD209 BV421 | 9E9A8 | Biolegend | 330117 | B368886 | 1:40 |
| Live/Dead Aqua | N/A | Thermo | L34957 | 2204201 | 1:500 |
